# Tunable gene expression in zebrafish using RiboSCALE

**DOI:** 10.64898/2026.05.08.723891

**Authors:** Thomas P. Rynes, Eiman A. Osman, Maureen McKeague, Karen Mruk

**Affiliations:** Department of Pharmacology and Toxicology, Brody School of Medicine, East Carolina University, Greenville, NC, USA; Department of Chemistry, Faculty of Science, McGill University, Montreal, QC H3A 0B8, Canada; Department of Pharmacology and Therapeutics, Faculty of Medicine and Health Sciences, McGill University, Montreal, QC H3G 1Y6, Canada

**Author notes:** Address correspondence to: Dr. Karen Mruk, Department of Pharmacology and Toxicology, East Carolina University, Brody School of Medicine, 600 Moye Boulevard, BSOM 6s20, Greenville, NC 27858.

**Keywords:** Chemogenetics, ribozyme, aptamer, zebrafish

## Abstract

Chemogenetic tools enable conditional control of gene expression during embryonic development and regeneration. However, many conditional tools induce constitutive or one-way activity precluding temporal resolution of gene function or require the use of multiple transgenic lines. We developed an RNA-based chemogenetic approach to induce gene expression in zebrafish embryos and larvae. We demonstrate that a gene of interest can be turned on in a time-dependent and concentration-dependent manner. Using this approach, we have characterized two different aptamers for future investigation.

## INTRODUCTION

Tissue development, aging, and regeneration rely on signaling networks composed of multiple genes. To interrogate these pathways, transient expression of exogenous DNA or mRNA has been used to overexpress a gene of interest. Although transient genetic approaches are quick and require little animal facility space, both approaches suffer from a short lifespan and lack of temporal and spatial control. As such, several transgenic technologies have been developed to express genes of interest that can last throughout the lifetime of the organism. For example, heat shock elements have been used to express genes in *Xenopus* (Bienz and Pelham, 1986) and mice (Chen et al., 2024), mosaically label *Drosophila* embryos (Halfon et al., 1997), rescue developmental defects in *C. elegans* (Bacaj and Shaham, 2007), and zebrafish embryos (Hao et al., 2013), promote regeneration after injury in adult zebrafish (Mokalled et al., 2016, Mukherjee et al., 2021) and human cells (Rome et al., 2005). However, heat shock promoters can vary among different animals, making optimization of these tools between species challenging (Astakhova et al., 2015). Given this limitation, the powerful Gal4/UAS (Scheer and Campos-Ortega, 1999, Aigaki et al., 2001, Hirsch et al., 2002, Halpern et al., 2008, Iacopino et al., 2022) and Cre-lox (Sauer, 1998, Mosimann and Zon, 2011, Hubbard, 2014) systems are still the most widely used technologies for targeted gene expression in model organisms. However, these systems also have limitations including the need for multiple transgenic lines, constitutive activity of transcriptional regulators or recombinases, reduced temporal control in some applications, and potential toxicity depending on the experimental context.

To get around these limitations, technologies that can be induced with small molecules have been developed in various systems. These chemogenetic technologies primarily use one of two approaches: utilizing a chemically-inducible promoter (Hagihara et al., 1999, Lottmann et al., 2001, Buchholz, 2012) or developing chimeric proteins (Osterwalder et al., 2001, Esengil et al., 2007, McClure et al., 2022) that are activated by a small molecule. However, these approaches still often require >1 transgenic line. Further, the small molecule selection for these technologies is limited, precluding the widespread use of these techniques for multiple biological questions. For example, a tetracycline-inducible promoter may have limited utility in infection studies where antibiotics may be used as an experimental group. Similarly, a chimeric protein utilizing the estrogen-receptor is not practical for studies focused on estrogen signaling. One approach for expanding the chemogenetic toolkit is the use of aptamers. Aptamers are structured nucleic acids that bind target molecules through three-dimensional, shape-dependent interactions with high specificity and affinity (DeRosa et al., 2023). Because aptamers can be generated against a broad range of targets including small molecules, ions, metabolites, and proteins, they provide a versatile platform for engineering chemically responsive systems in biological models (Townshend et al., 2021). In particular, aptamers can be coupled to ribozymes to create RNA switches that regulate gene expression in response to small molecule binding (Tarnowski and Gorochowski, 2022).

In this report, we investigate the ability of RNA aptamers fused to a ribozyme to control gene expression using two distinct molecules. We demonstrate that our previously reported zebrafish-compatible ribozyme platform incorporating the theophylline aptamer can induce gene expression in mammalian cells, zebrafish embryos, and zebrafish larvae in a dose-dependent manner. Using the ubiquitin promoter, we show that this technology can be applied in multiple tissue types. Finally, we demonstrate that our screening platform in mammalian cells directly translates into stable line generation in zebrafish, bypassing the need for optimization using transient approaches. We anticipate that aptamer-based control of gene expression will be a versatile approach for studying developmental, aging, and regenerative biology.

## RESULTS

### Optimization and characterization of RiboSCALE, a chemogenetic gene expression system

We previously showed that a ribozyme-aptamer system could function in zebrafish embryos as a biosensor for theophylline (Osman et al., 2024). Therefore, we first sought to test whether this system could work in a tunable way to control gene expression. To facilitate screening between zebrafish and mammalian cells, we used the bidirectional zebrafish pDIVE-Tol2 plasmid (*ubb:GFP-theo; ubb:mCherry*) (Osman et al., 2024). In this construct, GFP expression is controlled by a hammerhead ribozyme fused to a theophylline aptamer while mCherry is constitutively expressed. We transfected HEK293 cells with the construct and treated cells with different concentrations of theophylline for 24 or 48 hours (Fig. 1). Quantification of fluorescence from the GFP and mCherry channels led to a 2.5–2.8-fold change in the GFP/mCherry ratio with 24 hours of treatment. A larger increase was observed in cells incubated in theophylline for longer (48 hours, 3.2–3.8-fold change).

**Figure 1.**
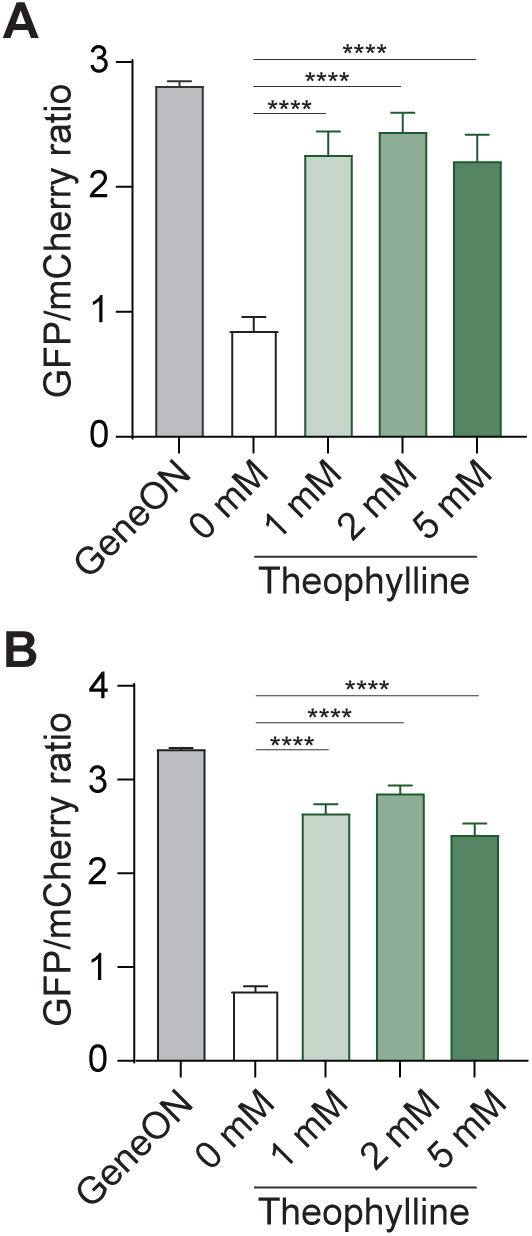
The pDIVE-Tol2 plasmid containing a theophylline aptamer controls GFP expression in HEK293 cells. Cells were transfected with the (*ubb:GFP-ON; ubb:mCherry*) or (*ubb:GFP-theo; ubb:mCherry*) plasmid and immediately treated with different concentrations of theophylline for (A) 24 h or (B) 48 h. Quantification shows the mean GFP/mCherry ratio ± sd. n=3 biological replicates for each condition. *p* values from a one-way ANOVA with Dunnett’s multiple comparisons test:: p<0.0001.

To characterize the utility of this system in a model organism, we generated zebrafish carrying a ribozyme-theophylline aptamer construct under control of the ubiquitin (*ubb*) promoter (Mosimann et al., 2011). Multiple founders were screened for single insertions of the expression construct and raised to generate stable lines. When crossed to WT AB zebrafish, both lines yielded embryos that exhibited theophylline-inducible GFP expression (Fig. S1). One of the heterozygous lines exhibited more robust switching throughout the embryo and we used that line for the rest of the experiments.

We previously showed that 1 mM theophylline did not cause developmental defects in early embryos but that embryos exposed to 2 mM exhibited developmental phenotypes when exposed for longer than 8 hours (Osman et al., 2024). Therefore, we dosed embryos at 24 hours post fertilization (hpf) in either 1 mM, 2 mM, or 5 mM theophylline and quantified pixel intensity after shorter incubation times (2–8 h) (Fig. 2). Quantification of pixel intensity from the trunk of the embryos two hours after dosing showed no statistical increase in the GFP/mCherry ratio. In contrast, embryos in all three concentrations of theophylline began to show an increase in the GFP/mCherry ratio after four hours of incubation with theophylline (1.4–2-fold change). After eight hours, embryos in 1 mM theophylline had a 2.8-fold change in the GFP/mCherry ratio whereas embryos in 5 mM theophylline had nearly a 5-fold change in the GFP/mCherry ratio (Fig. 2). Further, the increase in GFP expression in just eight hours was similar to the increase in GFP expression after incubation in 1 mM theophylline for 24 hours (Fig. S1), indicating that the amount of GFP overexpression can be manipulated with the length of time exposed to theophylline or the concentration of theophylline. As such, we named this new method ribozyme-Small-molecule Controlled Adjustable Level Expression (riboSCALE).

**Figure 2.**
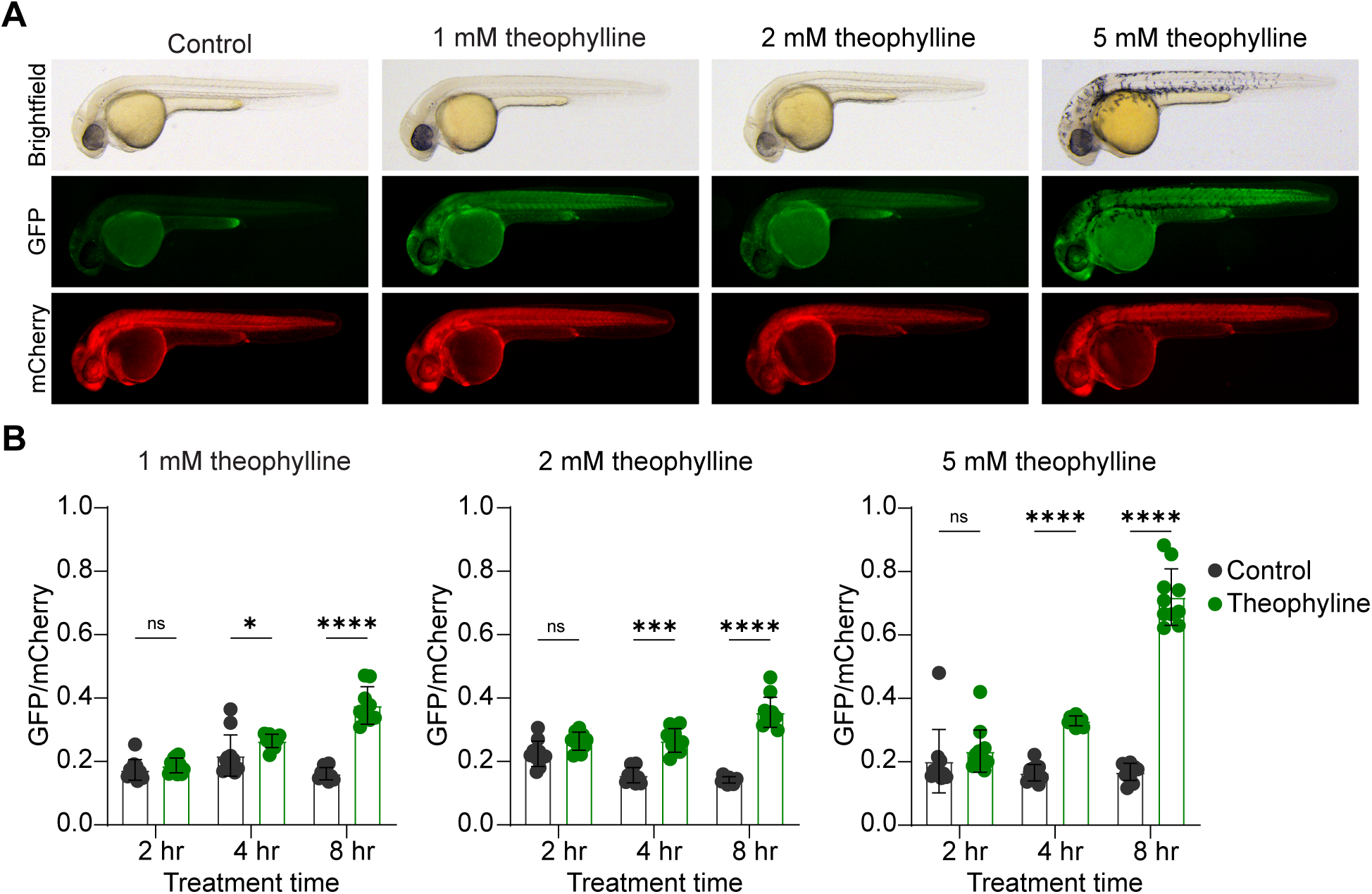
Theophylline induces GFP expression in *Tg(ubb:GFP-theo; ubb:mCherry)* embryos in a time and dose-dependent manner. (A) Representative bright-field and epifluorescence micrographs from 24 hpf *Tg(ubb:GFP-theo; ubb:mCherry)* embryos treated with theophylline for 8 h. Theophylline induces GFP expression in a concentration-dependent manner. Embryo orientation: lateral view, anterior left. Scale bar: 500 µm. (B) Quantification of GFP and mCherry fluorescence in trunk of *Tg(ubb:GFP-theo; ubb:mCherry)* embryos and sibling controls treated with theophylline. The average GFP/mCherry ratio ± sd is graphed with individual fish plotted. Adjusted p-values from two-way ANOVA with Šídák’s multiple comparisons test: *p=0.0314, ***p=0.0003, ****p<0.0001. n=10-12 embryos per condition

### Generalizability of RiboSCALE

Many tools are optimized in early embryos but then show reduced efficacy in specific tissues. To first test whether the ribozyme would function in multiple tissues, we generated transgenic fish containing an inactive ribozyme. These GeneON fish were imaged through the brain, heart, and trunk at 48 and 72 hpf (Fig. S2). Quantification of the GFP/mCherry pixel intensity showed no difference in GFP expression within the heart, neurons, or somites within a single age (Fig. S2A), indicating the construct was expressed in all tissues. Further, the GFP/mCherry ratio increased with embryo age (Fig. S2B), suggesting that as tissues develop they continuously express the transgene.

We next tested zebrafish expressing the theophylline-responsive ribozyme-aptamer construct (Fig. 3). Incubating 24 hpf embryos in 1 mM theophylline for 24 hours led to robust GFP expression leading to an 8.2-fold increase in the GFP/mCherry ratio in the heart, 6.2-fold increase in neurons, and 7.6-fold increase in somites (Fig. 3A). Unexpectedly, when dosing embryos with 1 mM theophylline at 48 hpf, we found an increase in the GFP expression in the heart (12.6-fold) but decrease in somites (5.9-fold, Fig. 3B). To determine whether the decrease in somites may be due to toxicity of theophylline in older embryos, we repeated the toxicity study on older embryos (Fig. S3). We found that 48 hpf embryos incubated in 1 mM and 2 mM theophylline were shorter than embryos not exposed to theophylline. In addition, 50% of 72 hpf embryos incubated in 1 mM failed to develop a swim bladder, indicating that prolonged exposure to theophylline may cause developmental toxicity, which may reduce GFP expression.

**Figure 3.**
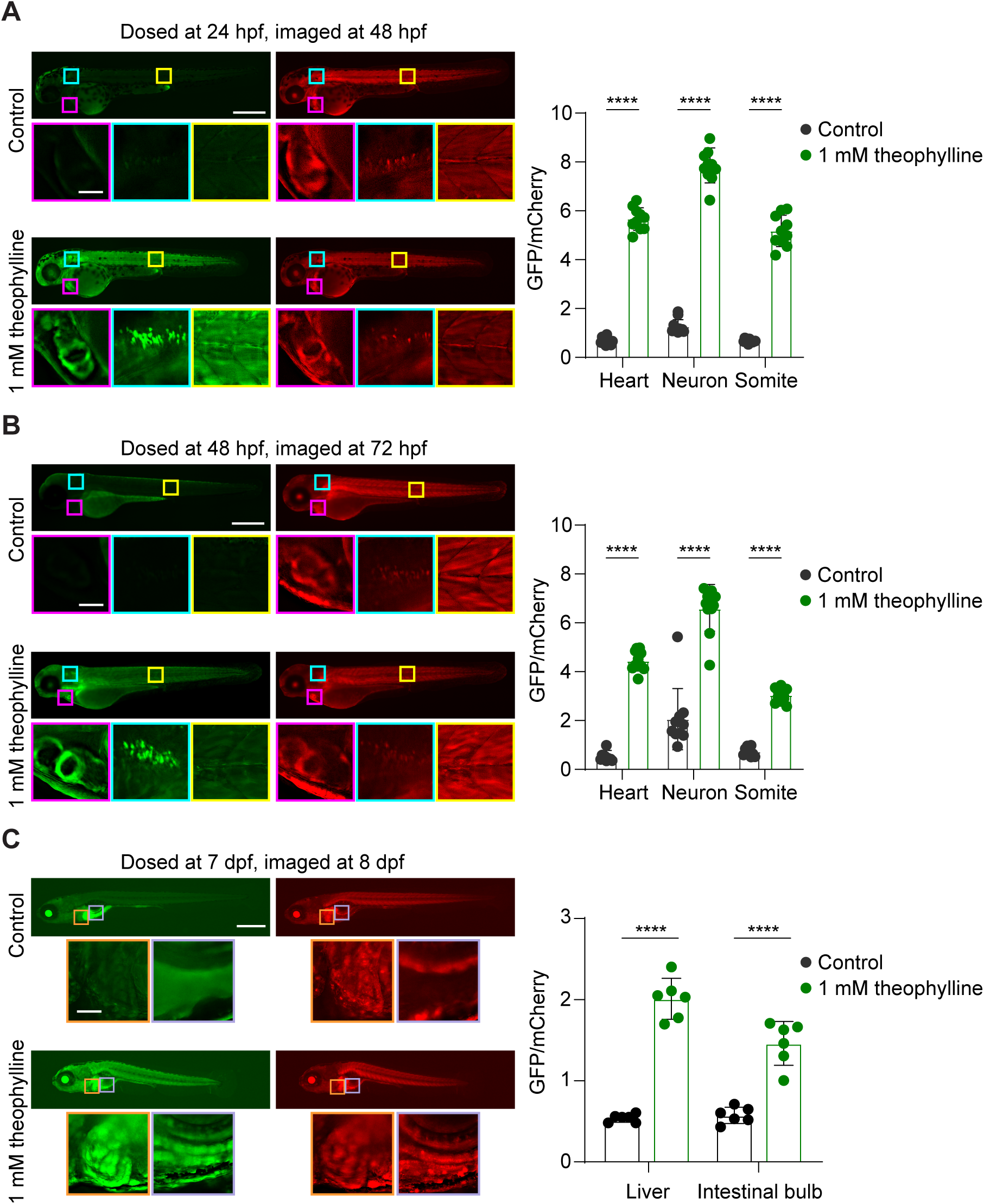
Theophylline induces GFP expression in *Tg(ubb:GFP-theo; ubb:mCherry)* embryos and larvae in multiple tissue types. Left: Representative epifluorescence micrographs from *Tg(ubb:GFP-theo; ubb:mCherry)* embryos dosed at (A) 24 hpf, (B) 48 hpf, and (C) 7 dpf for 24 h. Theophylline induces GFP expression in neurons, somites, heart, liver, and gut cells. Embryo and larvae orientation: lateral view, anterior left. Scale bar: 500 µm, inset: 50 µm. Right: Quantification of GFP and mCherry fluorescence *Tg(ubb:GFP-theo; ubb:mCherry)* embryos and larvae. The average GFP/mCherry ratio ± sd is graphed with individual fish plotted. Adjusted p-value from two-way ANOVA with Šídák’s multiple comparisons test: ****p<0.0001. n=10-12 embryos and 6 larvae per condition

We reasoned that some toxicity may be due to the inability of an embryo to metabolize theophylline as the liver has not yet developed (Wilkins and Pack, 2013). Therefore, we also treated larvae at 5 and 7 days post fertilization (dpf) with 1 mM theophylline for 24 hours. Indeed, larvae incubated with 1 mM theophylline at 5 dpf, developed similarly to control larvae, suggesting some toxicity could be mitigated by hepatic metabolism of theophylline (Fig. S3). Quantification of the GFP/mCherry ratio in 7 dpf larvae showed robust GFP expression in the liver (3.7-fold change) and cells lining the intestinal bulb (2.5-fold change) (Fig. 3C). Taken together, the riboSCALE system can be utilized to overexpress a gene in multiple ages and tissues.

### Expanding the RiboSCALE toolkit

Two limitations in current chemogenetic technologies for animal models are the potential toxicity of some commonly used molecules and a lack of technologies that utilize novel small molecules. Further, screening thousands of different small molecule ribozyme-aptamer pairs in an animal is challenging. Therefore, we next sought to determine whether the mammalian cell screening platform could be used to evaluate alternative ribozyme-aptamer pairs responsive to potentially less toxic small molecules prior to generating stable transgenic zebrafish lines. Specifically, we transfected HEK293 cells with our Tol2-bidirectional plasmid which had GFP under control of a ribozyme-folinic acid aptamer (*ubb:GFP-folinic; ubb:mCherry*). We reasoned that since folinic acid is used in patients to counteract chemotherapy toxicity (1995, 1999), it would be less toxic in zebrafish. We treated cells with different concentrations of folinic acid for 24 or 48 hours (Fig. 4). Quantification of pixel intensity from the GFP and mCherry channels led to a 1.1–1.4-fold change in the GFP/mCherry ratio in treated cells, indicating that although the folinic acid aptamer showed lower gene activation than the theophylline aptamer, it was still able to induce gene expression.

**Figure 4.**
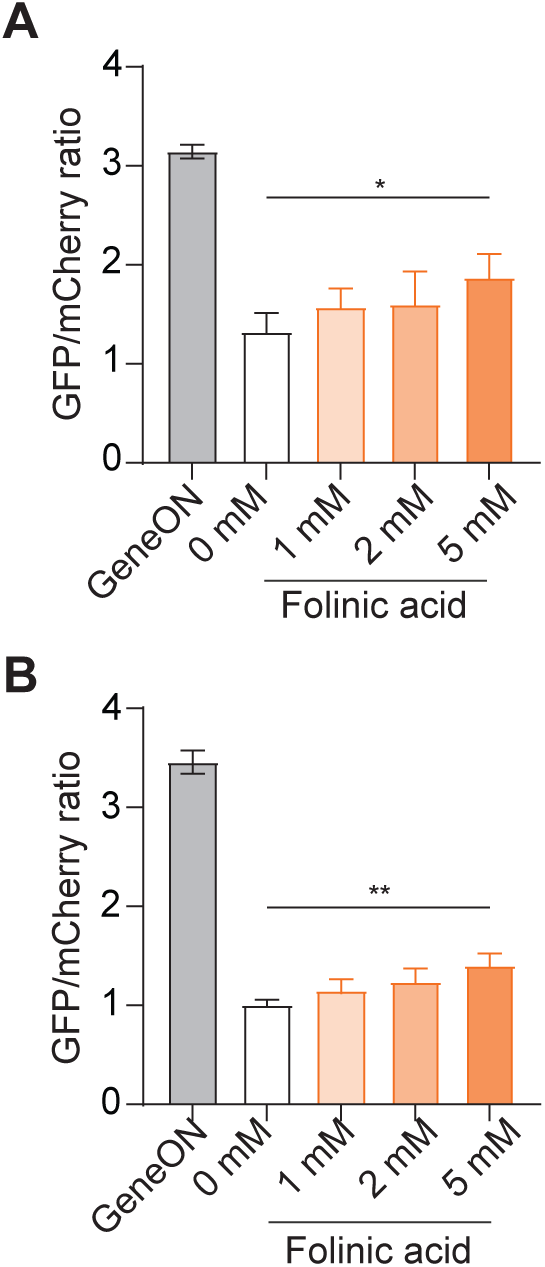
The pDIVE-Tol2 plasmid containing a folinic acid aptamer controls GFP expression in HEK293 cells. Cells were transfected with the (*ubb:GFP-ON; ubb:mCherry*) or (*ubb:GFP-folinic; ubb:mCherry*) plasmid and immediately treated with different concentrations of theophylline for (A) 24 h or (B) 48 h. Quantification shows the mean GFP/mCherry ratio ± sd. n=3 biological replicates for each condition. *p* values from a one-way ANOVA with Dunnett’s multiple comparison test: *p=0.0357, **p=0.0039

To determine whether this would translate to zebrafish, we generated transgenic zebrafish expressing the folinic aptamer, *Tg(ubb:GFP-folinic; ubb:mCherry),* and treated 24 hpf embryos with 1 mM, 2 mM, or 5 mM folinic acid as described for the theophylline aptamer. Consistent with mammalian cell data, we found that the ribozyme-folinic acid aptamer only increased the GFP/mCherry ratio in the trunk of the embryo 1.1–1.2-fold (Fig. 5). To ensure the reduced switching was not due to toxicity, we repeated the toxicity experiments by incubating embryos at 24 hpf, 48 hpf, and 72 hpf. Embryos of all three ages showed no change in embryo length compared to control embryos, suggesting that the reduced GFP expression is not due to toxicity of folinic acid (Fig. S4). Quantification of the GFP/mCherry ratio in the heart and somites showed no statistical change in GFP expression, whereas there was an increase in neurons in embryos treated at 24 hpf but not at 48 hpf (Fig. 6). This is consistent with the ability of folinic acid to support brain development by bypassing transport systems (Frye et al., 2020). Given the developmental differences in folinic acid transport and uptake, it is perhaps unsurprising that this molecule only induced expression in young embryos. Taken together, our data suggest that ribozyme-aptamer screening in mammalian cells can directly translate to transgenic zebrafish lines.

**Figure 5.**
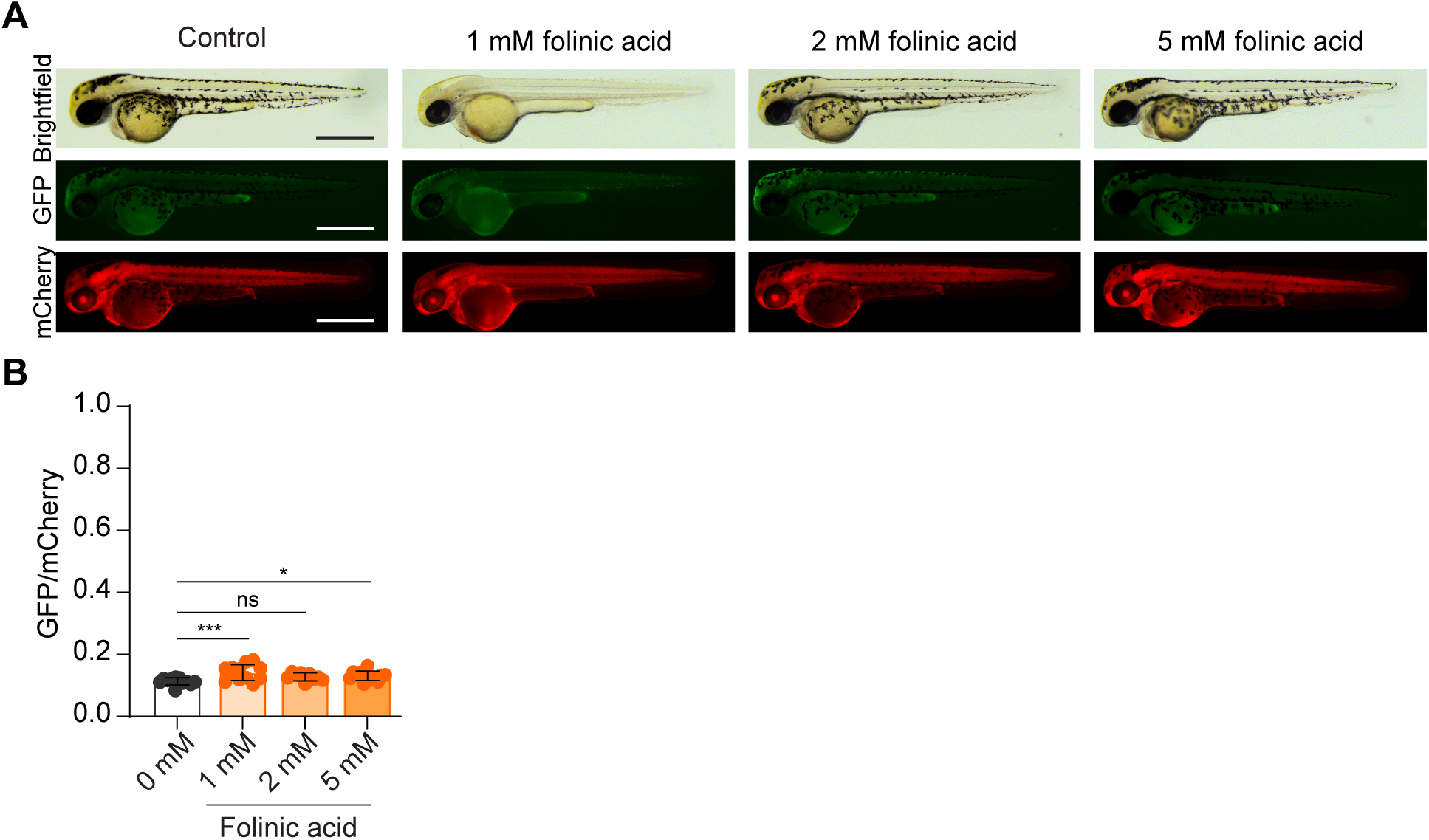
Folinic acid induces GFP expression in *Tg(ubb:GFP-folinic; ubb:mCherry)* embryos in a time and dose-dependent manner. (A) Representative bright-field and epifluorescence micrographs from 24 hpf *Tg(ubb:GFP-theo; ubb:mCherry)* embryos treated with folinic acid for 24 h. Folinic acid induces weak GFP expression. Embryo orientation: lateral view, anterior left. Scale bar: 500 µm. (B) Quantification of GFP and mCherry fluorescence in trunk of *Tg(ubb:GFP-folinic; ubb:mCherry)* embryos treated with folinic aicd. The average GFP/mCherry ratio ± sd is graphed with individual fish plotted. Adjusted p-values from one-way ANOVA with Dunnett’s multiple comparisons test: *p=0.0445, ***p=0.0008 n=10-12 embryos per condition

**Figure 6.**
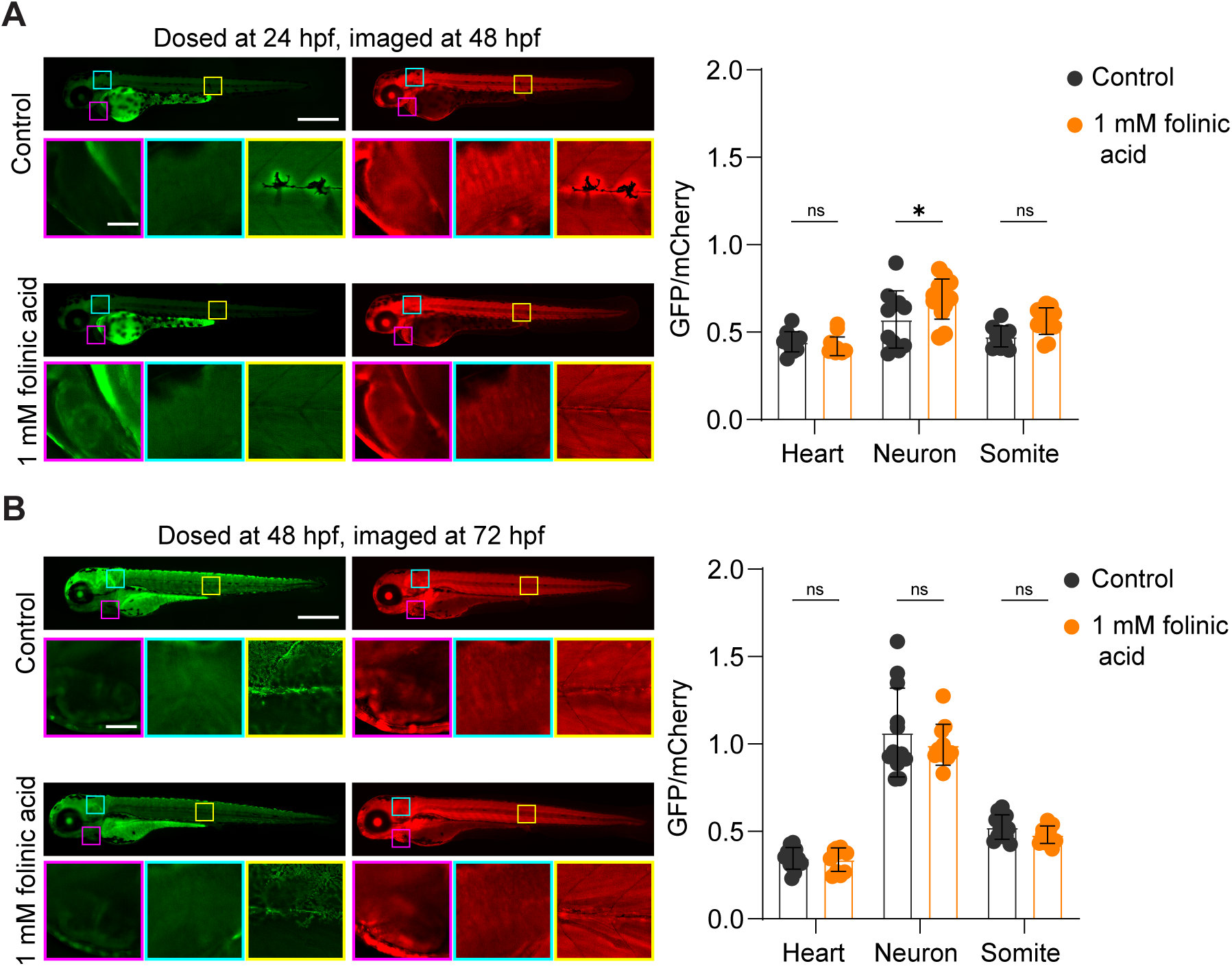
Folinic acid induces GFP expression in neurons of 24 hpf *Tg(ubb:GFP-folinic; ubb:mCherry)* embryos. Left: Representative epifluorescence micrographs from (A) 24 hpf and (B) 48 hpf *Tg(ubb:GFP-theo; ubb:mCherry)* embryos treated with 1 mM folinic acid for 24 h. Folinic acid induces weak GFP expression. Embryo orientation: lateral view, anterior left. Scale bar: 500 µm, inset: 50 µm. Right: Quantification of GFP and mCherry fluorescence *Tg(ubb:GFP-folinic; ubb:mCherry)* embryos treated with folinic aicd. The average GFP/mCherry ratio ± sd is graphed with individual fish plotted. Adjusted p-values from one-way ANOVA with Dunnett’s multiple comparisons test: *p=0.0445, ***p=0.0008 n=10-12 embryos per condition

## DISCUSSION

Our results establish that ribozyme-aptamer systems can be used to control expression of a gene of interest in both mammalian cells and zebrafish with tunable control. Our findings complement previous work demonstrating that RNA aptamers can work in invertebrate animal models (Wurmthaler et al., 2019, Fang et al., 2023) and plants (Shanidze et al., 2020). Our direct comparison between transient transfection in mammalian cells and stable transgenic lines in zebrafish highlights the species-independent use of this technology. The approach we described allows follow-up screens in which gene expression can be affected by different molecules. These attributes highlight the versatility of the ribozyme-aptamer system as a chemogenetic tool. The functionality of this tool could be further enhanced by using alternative *cis-*regulatory sequences to restrict expression to specific cell types. In addition, the concentration-dependent and time-dependent induction observed in both mammalian cells and zebrafish suggests that these systems may provide a useful framework for studying dynamic biological processes that require graded or reversible control of gene expression.

Although the theophylline aptamer induced robust gene expression, induction with the folinic acid aptamer was significantly less. This could be due to limitations in tissue permeability or affinity for the aptamer. Given there are approximately 350,000 different chemical substances registered for commercial use, both limitations can be overcome by screening new aptamer-molecule pairs. Our data show that screening aptamers in mammalian cells using the pDIVE-Tol2 plasmid reports on efficacy of a switch in zebrafish, increasing the potential for discovering new pairs that can directly translate into stable transgenic models without the need for testing using transient approaches such as RNA injection (LaBelle et al., 2021, Mruk et al., 2020). Further, although the folinic acid aptamer exhibited reduced activation relative to the theophylline aptamer, folinic acid exposure caused substantially less developmental toxicity, highlighting the potential utility of alternative small molecules for long-term or late-stage studies. However, these findings also highlight the importance of balancing activation efficiency with ligand toxicity, tissue permeability, and developmental stage when selecting ribozyme-aptamer pairs for in vivo applications.

As research in aging and regenerative medicine continues to grow, there is a growing need to develop new tools for conditional overexpression and genetic knockdown in older animals. A recent study in zebrafish embryos showed that the T3H48 ribozyme can act as a conditional knockdown of a gene of interest when introduced to the intron of an endogenous gene of interest (Juan et al., 2025). Unlike the ribozyme-aptamer systems used here, which are derived from the satellite Tobacco Ringspot Virus (sTRSV) hammerhead ribozyme scaffold and optimized for ligand-dependent modulation of gene expression, the engineered T3H48 hammerhead ribozyme was developed for highly efficient constitutive cleavage and conditional knockdown of endogenous genes. While the engineered T3H48 ribozyme was optimized for maximal cleavage efficiency (Chi et al., 2008), the sTRSV-derived ribozymes used in this study retain the dynamic range necessary for tunable induction in response to small molecules. Together, these studies highlight the versatility of engineered ribozymes for vertebrate gene regulation and open the possibility of combining different aptamers with distinct ribozyme architectures to control temporal dynamics, multiplex regulation of >1 gene, or model diseases where gene expression changes with aging. Future studies aimed at identifying additional aptamer-molecule pairs with improved activation profiles and reduced toxicity may further expand the utility of these systems across developmental and regenerative contexts. Moreover, given that aptamers are small (<100 bp), they are amenable to genome insertion through TALEN- or CRISPR-mediated techniques (Auer and Del Bene, 2014, Auer et al., 2014). Based on these capabilities, we anticipate that ribozyme-aptamer systems will be versatile tools for deconstructing developmental and regenerative mechanisms in whole organisms.

## MATERIALS AND METHODS

### Small molecules

Theophylline was purchased from Millipore Sigma (#T1633). (6R,S)-5-formyl-5,6,7,8-tetrahydrofolic acid calcium salt (#16.220-5) was purchased from Schircks Laboratories (Bauma, Switzerland) or Millipore Sigma (#47612). Gibco Dulbecco’s Modified Eagle Medium (DMEM), antibiotic–antimycotic (100X), trypsin, fetal bovine serum (FBS, #A3160702), phosphate buffered saline (PBS), and Lipofectamine 3000 (#L3000015) were purchased from Fisher Scientific.

### pDIVE-Tol2 plasmids

The inactive ribozyme and aptamer plasmids were previously reported (Osman et al., 2024). Briefly these constructs were generated by cloning RNA switch inserts, produced via overlap extension PCR, into a zebrafish-compatible backbone at the XmaI site within the 3′ untranslated region (UTR) of GFP using Gibson assembly. Plasmids were propagated in chemically competent TOP10 *Escherichia coli*, purified, and sequence-verified by Sanger sequencing.

### Cell culture and transfection

Human embryonic kidney (HEK293T) cells were cultured in DMEM supplemented with 10% FBS and 1% antibiotic–antimycotic (penicillin–streptomycin). Cells were maintained at 37 °C in a humidified atmosphere with 5% CO₂. HEK293T cells were routinely tested for mycoplasma contamination and were not used beyond passage 25. Cells were seeded at 2 × 10⁵ cells per well in a 24-well plate. After 24 h, cells were transfected with 0.5 µg of control or RNA switch plasmids using Lipofectamine 3000 and immediately treated with theophylline or folinic acid at concentrations of 0, 1, 2, and 5 mM. Cells were then incubated for an additional 24 or 48 h.

### Measuring RNA switch activity in mammalian cells

Following 24 or 48 h incubation, media was removed and cells were lysed by adding Glolysis buffer (150 µL) to each well, followed by a 5 min incubation at room temperature. Lysates were transferred directly to a 96-well black plate. GFP (ex/em: 488/510 nm) and mCherry (ex/em: 587/610 nm) were measured on a BioTek Cytation 5 plate reader. Ratios of GFP/mCherry were calculated and analyzed using GraphPad Prism (v8.4.3). Each experiment included technical triplicates and was repeated three times to generate biological triplicates. Data were plotted as mean ± standard deviation.

### Zebrafish husbandry

Zebrafish [WT AB (ZL1) and Casper (ZL1714)] were purchased from the Zebrafish International Resource Center (ZIRC). Adult fish were maintained at 28.5°C in a recirculating system (Iwaki Aquatic). A 14:10 hour light:dark cycle was used, and fish were fed twice daily. Morning diet of brine shrimp (Brine Shrimp Direct), and afternoon feeding of Zeigler adult zebrafish diet (Pentair Aquatic Ecosystems). Embryos were obtained through normal mating practices and stored at 28.5°C in 1x E3 buffer. Zebrafish larvae were raised to 5 days post fertilization (dpf) and then fed rotifers (Reed mariculture) from 5 dpf to approximately 14 dpf. From 14 dpf to sexual maturity fish were fed a brine shrimp diet. Zeigler diet was added to the standard schedule once larvae were old enough to ingest the pellet. The IACUC committee at East Carolina University, Greenville, NC, USA approved all animal procedures (AUP#W262).

### Zebrafish transgenic lines

To generate a stable transgenic ribozyme-aptamer lines, plasmid DNA (25-50 pg) and *Tol2* mRNA (50 pg) were premixed and injected into one-cell-stage WT AB embryos. Fish were raised to 24 hpf and checked for successful injection using the mCherry channel. Positive expression fish were raised to adulthood as described above. The F0 adults were mated with Casper fish to identify founders with germline transmission. For all experiments, at least two different F0s that yielded F2 generations with monoallelic expression were used to establish transgenic lines.

### Zebrafish imaging

Live zebrafish were mounted laterally on a glass depression slide using 1.5% low melting point agarose. For dosing and toxicity studies, fish were imaged on a Leica M165 FC equipped with a Planapo 1.0x M-series objective and K7 camera or an Olympus SZX16 equipped with a DP80 camera and SDF PLANO 1XPF objective. For tissue-specific experiments, fish were imaged on a Leica DM6 with K8 CMOS camera and a 20x/NA 0.5 water-immersion objective. Z-stacks were generated from images taken at 5 μm intervals. Images were saved and imported into FIJI (Schindelin et al., 2012) for all analysis.

### Toxicity assays

WT fish were dosed with 0.5 mM, 1 mM, and 2 mM of drug (folinic acid or theophylline) directly dissolved in E3 buffer. Four ages of fish were tested (24 hpf, 48 hpf, 72 hpf, and 5 dpf) and tracked during development for a 24-hour period. Fish were imaged as described above using the bright filed channel at 2, 4, 8, and 24 h of drug exposure. The overall length of the fish was measured using FIJI or Olympus CellSense software and recorded in an excel spreadsheet.

### Fluorescence intensity quantification for zebrafish

Fluorescence intensity quantification was preformed using FIJI. Pixel intensity was determined by measuring the minimum, maximum and mean grey value in a region of interest (ROI) on the image in both the GFP and mCherry channels. The ROI was drawn in FIJI with either the rectangle or polygon tool. The ROI for each age of fish and for each experiment were saved to maintain consistency of the area. Two measurements were taken, one with the ROI over the desired location (heart, neurons, somites, etc) and one measurement over the background of the image. The two measurements were then subtracted from each other to produce a “true” grey value measurement. All measurements were saved in an excel spread sheet. The ratio of GFP/mCherry intensity was calculated by dividing the “true” value of the GFP by the “true” value of the mCherry. All imaged zebrafish that had mCherry fluorescence were used in the quantification with no exclusions.

### Statistics of zebrafish experiments

For zebrafish experiments, at least two breeding tanks, each containing 2-3 males and 3-5 females from separate stocks, were set up to generate embryos. Embryos from each tank were randomly distributed across tested conditions. Unfertilized eggs and developmentally abnormal embryos were removed prior to imaging or compound treatment. No statistical methods were used to determine sample size per condition. Values for individual fish are plotted, and each distribution was assessed using the Shapiro-Wilk test for normality. All graphs were prepared in GraphPad Prism and statistical testing using was completed in the software (Dotmatics). Individual tests are reported in the figure legends.

## ACKNOWLEDGMENTS

We gratefully acknowledge financial support from NIH (R21GM143565 to M.M. and K.M, R03NS136719 to K.M.) and the Natural Sciences and Engineering Research Council of Canada (RGPIN-2026-06146 to M.M).

## AUTHOR CONTRIBUTIONS

Conceptualization: MM, KM; Methodology: TPR, EAO, MM, KM; Validation: TPR, EAO, MM, KM; Formal Analysis: TPR, EAO, MM, KM; Investigation: TPR, EAO; Resources: TPR, EAO; Writing: TPR, EAO, MM, KM; Editing: TPR, EAO, MM, KM; Visualization: TPR, EAO, MM, KM; Supervision: MM, KM; Project Administration: KM; Funding acquisition: MM, KM

## SUPPORTING INFORMATION

Figs. S1-S3.

