## SupplementalPacket for "Tunable gene expression in zebrafish using RiboSCALE"

TABLE OF CONTENTS

Supplementary Figure 1 ……………………………………………………………………………..3

Supplementary Figure 2 ……………………………………………………………………………..4

Supplementary Figure 3 ……………………………………………………………………………. 5

### Figure S1

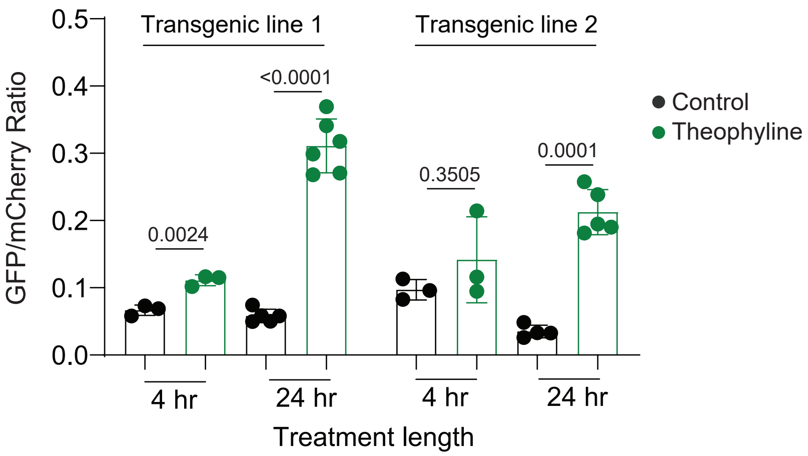


**Figure S1. Characterization of *Tg(ubb:GFP-theo; ubb:mCherry)* zebrafish.** Embryos from two independantly made *Tg(UbiGFP-theo; Ubi:mCherry)* lines were incubated in presence or absence of 1 mM theophylline and imaged after treatment. Quantification of pixel intensity of GFP and mCherry was calculated for the trunk of the zebrafish and the GFP/mCherry ratio plotted. Both lines show switching that increases with longer incubation times. The average GFP/mCherry ratio ± s.d. with values for individual fish shown. p values from a Kruskal-Wallist test with post-hoc Dunn’s test.

#### Figure S2

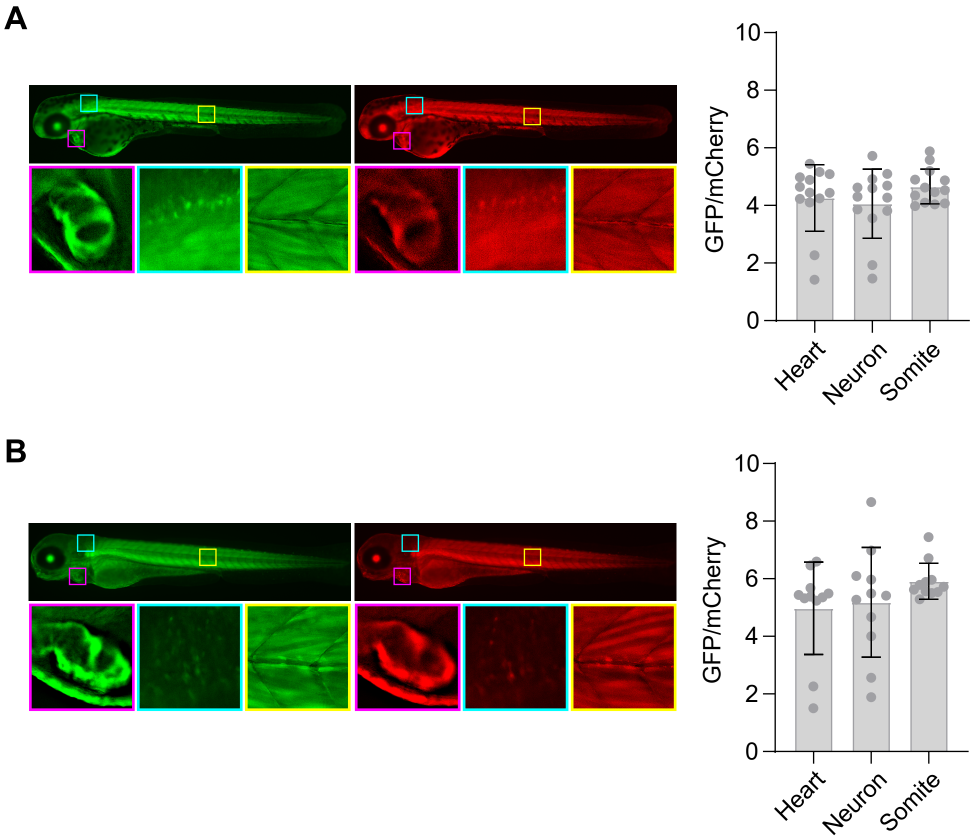


**Figure S2.** ***Tg(ubb:GFP-On; ubb:mCherry)* embryos have robust expression in different tissues during development.** Left: Representative micrographs from heterozygous embryos imaged at (A) 48 hpf or (B) 72 hpf. Right: Quantification of pixel intensity of the GFP and mCherry fluorescence in the trunk measured. The average GFP/mCherry ratio ± s.d. with values for individual fish shown. All embryos shown in lateral view, anterior left.

#### Figure S3

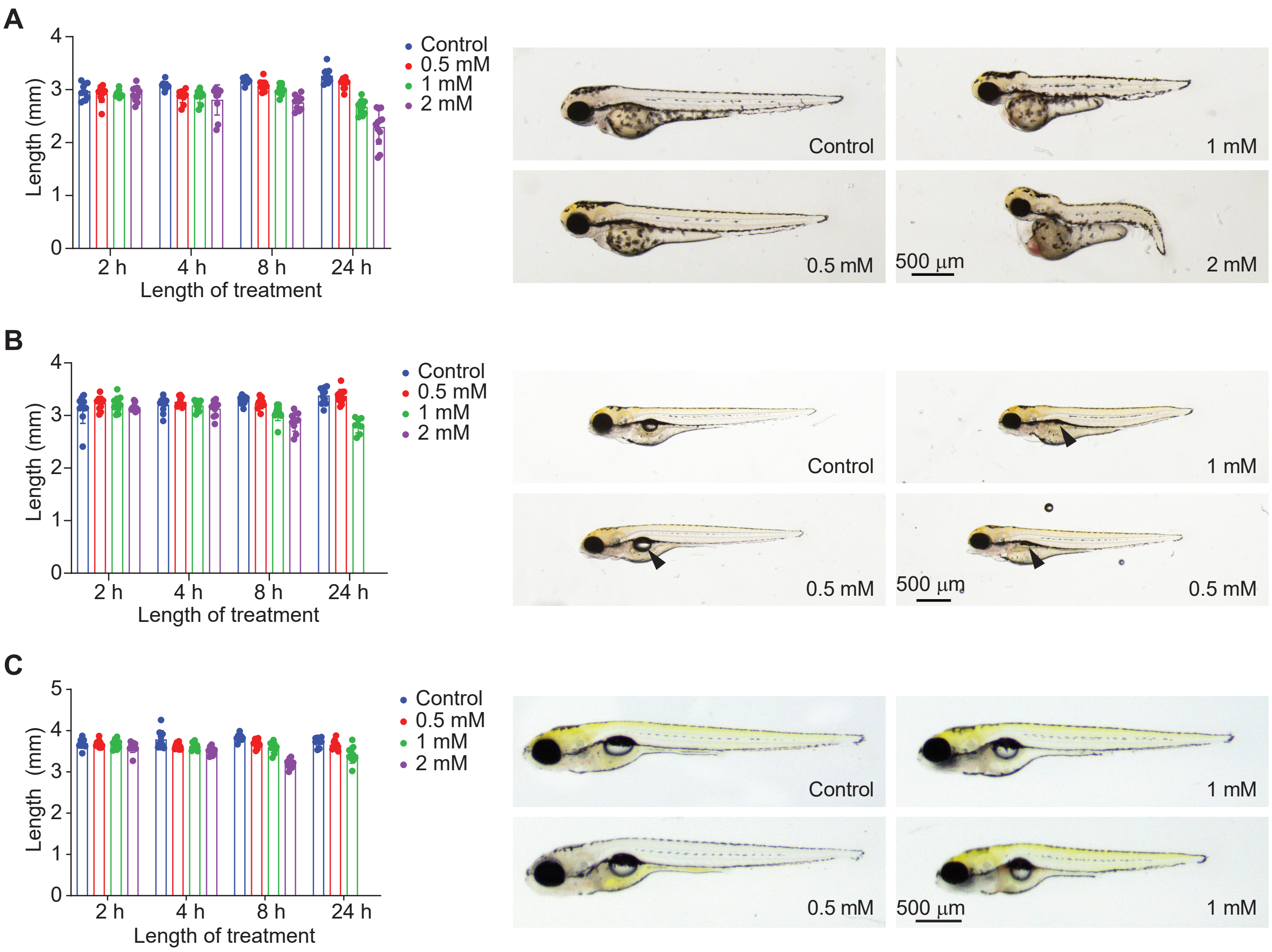


Fig. S3. Theophylline results in low toxicity to zebrafish embryos. WT embryos were exposed to various concentrations of theophylline at (A) 48 hpf, (B) 72 hpf, and (C) 5 dpf and imaged over time. Left: Quantification of total length with the average length ± s.d plotted. Individial fish are shown. Missing bars indicate embryos were too sick to image. Right: Representative micrographs after 24 h treatment. Micrographs are lateral view, anterior left. Treatment with theophylline can cause defects in swim bladder formation (arrowheads) and body length. n=10 embryos per condition

#### Figure S4

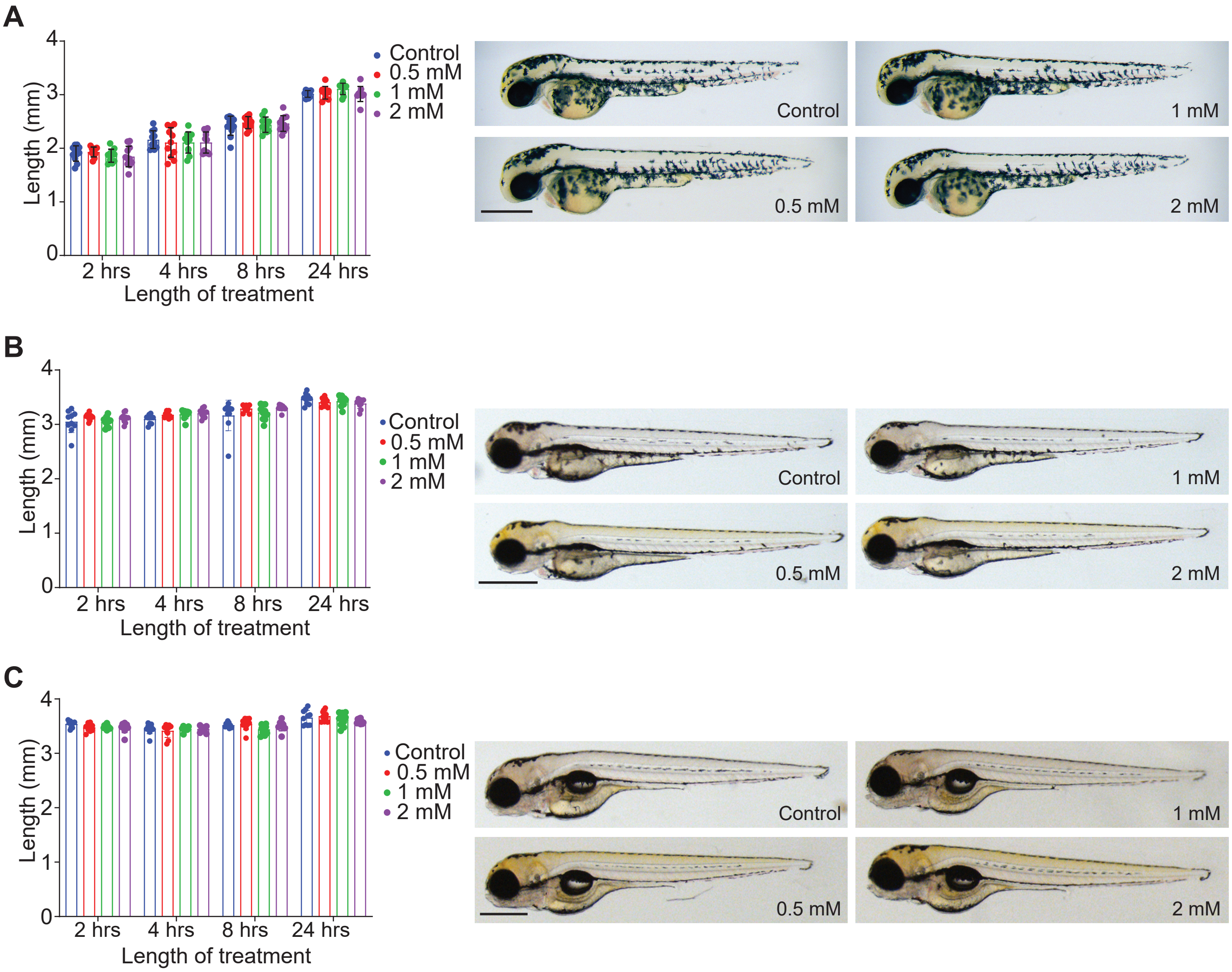


**Fig. S4. Folinic acid is non-toxic to zebrafish embryos.** WT embryos were exposed to various concentrations of folinic acid at (A) 24 hpf, (B) 48 hpf, and (C) 72 hpf and imaged over time. Left: Quantification of total length with the average length ± s.d plotted. Individial fish are shown. Right: Representative micrographs after 24 h treatment. Orientation for all micrographs: lateral view, anterior left. Scale bar: 500 μm
